## Supplementary material for "Time series-free rhythm profiling using COFE reveals multi-omic circadian rhythms in in vivo human cancers": SI Text, Tables and Figures

### Supporting Information: Text

#### S1 Text Theoretical results

##### A Alternative formulation of sparse cyclic PCA

In this section, we motivate sparse cyclic PCA from a parameter estimation perspective. Consider a simple generative model for a feature $j$ in the column-centered data matrix $\mathbf{X}$:

$$\mathbf{X}_{ij} = a_j \cos(\omega t_i) + b_j \sin(\omega t_i) + \varepsilon_{ij}, \quad \varepsilon_{ij} \sim \mathcal{N}(0, \sigma^2), \; i = 1, \dots N., \tag{S1}$$

where we have assumed that the noise variance is uniform across features for simplicity and $\omega = 2\pi/T$, where $T$ is the period of the rhythm. A *rhythmic* feature $j$ is defined by $(a_j, b_j) \neq (0,0)$ and an *arrhythmic* feature by

$$\mathbf{X}_{ij} = \varepsilon_{ij}, \quad \varepsilon_{ij} \sim \mathcal{N}(0, \sigma^2), \; i = 1, \dots N.$$

Eq. (S1) can be viewed as Fourier series approximation of a periodic signal at the fundamental frequency. The problem of determining the coefficients $(a_j, b_j)$ in the model (S1) is often called cosinor regression in the circadian literature [25]. An important caveat is that in our setting, the sample times $t_i$ are also unknown.

The log-likelihood of the data based on this generative model (S1) is:

$$L(\mathbf{X}) = \log \left[ \prod_{j=1}^{p} \frac{1}{(2\pi\sigma^2)^{N/2}} \exp\left( -\frac{\|\mathbf{X}_{.j} - a_j \mathbf{u}_1 - b_j \mathbf{u}_2\|^2}{2\sigma^2} \right) \right], \tag{S2}$$

where $\mathbf{u}_1 = [\cos(\omega t_1), \dots, \cos(\omega t_N)]^T$ and $\mathbf{u}_2 = [\sin(\omega t_1), \dots, \sin(\omega t_N)]^T$. By definition, the elements of $\mathbf{u}_1, \mathbf{u}_2$ satisfy the circular constraint, ${\mathbf{u}_1}_i^2 + {\mathbf{u}_2}_i^2 = 1$. Letting $\mathbf{v}_1 = [a_1, a_2, \dots, a_p]^T$ and $\mathbf{v}_2 = [b_1, b_2, \dots, b_p]^T$, the log-likelihood (S2) can be rewritten as:

$$L(\mathbf{X}) = -\frac{Np}{2} \log(2\pi\sigma^2) - \frac{\sum_{j=1}^{p} \|\mathbf{X}_{.j} - a_j \mathbf{u}_1 - b_j \mathbf{u}_2\|^2}{2\sigma^2} = -\frac{Np}{2} \log(2\pi\sigma^2) - \frac{\|\mathbf{X} - \mathbf{u}_1 \mathbf{v}_1^T - \mathbf{u}_2 \mathbf{v}_2^T\|_F^2}{2\sigma^2}. \tag{S3}$$

The maximum likelihood estimate for the model parameters $(a_j, b_j, t_i)$ is then equivalent to the rank-2 approximation introduced in (1)

$$\underset{\mathbf{u}_1, \mathbf{u}_2, \mathbf{v}_1, \mathbf{v}_2}{\operatorname{argmin}} \; \|\mathbf{X} - d\mathbf{u}_1 \mathbf{v}_1^T - d\mathbf{u}_2 \mathbf{v}_2^T\|_F^2, \;\; {\mathbf{u}_1}_i^2 + {\mathbf{u}_2}_i^2 = 1, \;\; \|\mathbf{v}_1\|_2 = \|\mathbf{v}_2\|_2 = 1,$$

where w.l.o.g. we have introduced a parameter $d$ to adapt the scale of the reconstruction after imposing an $l_2$-norm constraint on the $\mathbf{v}$s. To help identify the arrhythmic features, we encourage sparsity in $\mathbf{v}$ by enforcing a convex $l_1$-norm constraint ($\|\mathbf{v}_1\|_1 \leq s$, $\|\mathbf{v}_2\|_1 \leq s$). Ideally, we would like to enforce sparsity directly via a constraint on the $l_0$-norm, but since it is non-convex, following standard practice in the literature we relax it to an $l_1$-norm [47].

##### B Biconvexity of the optimization

Consider the optimized score in (3):

$$\max_{\mathbf{u}_1, \mathbf{v}_1, \mathbf{u}_1, \mathbf{v}_1, d} \left\{ 2d\mathbf{u}_1^T \mathbf{X} \mathbf{v}_1 + 2d\mathbf{u}_2^T \mathbf{X} \mathbf{v}_2 - d^2 N \right\}$$
$$\text{s.t. } \|\mathbf{v}_1\|_2 = \|\mathbf{v}_2\|_2 = 1, \|\mathbf{v}_1\|_1 \leq s, \|\mathbf{v}_2\|_1 \leq s, \;\; \mathbf{u}_{1i}^2 + \mathbf{u}_{2i}^2 = 1, \; \forall i \in \{1, \dots, N\}.$$

The objective function above is bilinear, i.e, linear in $\mathbf{u}$s for fixed $\mathbf{v}$s and vice versa. That is, the score can always be increased by scaling $\mathbf{u}$s and $\mathbf{v}$s by a factor larger than 1. Therefore, the solution to this problem remains unaltered when the constraints are expanded as follows:

$$\|\mathbf{v}_1\|_2 \leq 1,\ \|\mathbf{v}_2\|_2 \leq 1,\ \|\mathbf{v}_1\|_1 \leq s,\ \|\mathbf{v}_2\|_1 \leq s,\ \mathbf{u}_{1i}^2 + \mathbf{u}_{2i}^2 \leq 1,\ \forall i \in \{1,\ldots,N\}.$$

All the constraint sets are convex, since norms are convex and the intersection of convex sets is convex. Thus, convex constraint sets combined with a bilinear (biconvex) score proves the biconvexity of the optimization [46].

#### S2 Text Pseudocode of sparse cyclic PCA

**Algorithm 1** Sparse Cyclic PCA Algorithm

**Input:** $N \times p$ Data matrix $\mathbf{X}$, sparsity parameter $s$, tolerance $tol$, maximum iterations $max\_iter$
**Output:** $\mathbf{v}_1, \mathbf{v}_2$ (sparse loading vectors), $\mathbf{u}_1, \mathbf{u}_2$ (cyclic principal components), $d$ (scale factor)
Initialize $\mathbf{v}_1, \mathbf{v}_2$ with random draws of size $p \times 1$ from Laplace distribution
$\mathbf{v}_1 \leftarrow S(\mathbf{v}_1, \delta(s))$ // soft-threshold for sparsity constraint, Algorithm 2
$\mathbf{v}_2 \leftarrow S(\mathbf{v}_2, \delta(s))$ // soft-threshold for sparsity constraint, Algorithm 2
$\mathbf{v}_1 \leftarrow \mathbf{v}_1 / \|\mathbf{v}_1\|_2$ // $l_2$ normalize
$\mathbf{v}_2 \leftarrow \mathbf{v}_2 / \|\mathbf{v}_2\|_2$ // $l_2$ normalize
Initialize $\mathbf{u}_1 \leftarrow \cos(\phi)$ and $\mathbf{u}_2 \leftarrow \sin(\phi)$, where $\phi$ is random draw from uniform distribution $[0, 2\pi)$
$d \leftarrow \frac{(\mathbf{u}_1^T \mathbf{X} \mathbf{v}_1 + \mathbf{u}_2^T \mathbf{X} \mathbf{v}_2)}{N}$
$score \leftarrow 2d(\mathbf{u}_1^T \mathbf{X} \mathbf{v}_1 + \mathbf{u}_2^T X \mathbf{v}_2) - Nd^2$
$rss \leftarrow \|\mathbf{X}\|_F^2$
**repeat**
    // maximize **u**s keeping **v**s fixed
        $\mathbf{y}_1 \leftarrow \mathbf{X}\mathbf{v}_1$, $\mathbf{y}_2 \leftarrow \mathbf{X}\mathbf{v}_2$
        $\mathbf{y}_{circ} \leftarrow \sqrt{\mathbf{y}_1^2 + \mathbf{y}_2^2}$
        $\mathbf{u}_1 \leftarrow \mathbf{y}_1 / \mathbf{y}_{circ}$, $\mathbf{u}_2 \leftarrow \mathbf{y}_2 / \mathbf{y}_{circ}$
    // maximize **v**s keeping **u**s fixed
        $\mathbf{v}_1 \leftarrow S(X^T \mathbf{u}_1, \delta(s)) / \|S(X^T \mathbf{u}_1, \delta(s))\|_2$ // soft-threshold and normalize, Algorithm 2
        $\mathbf{v}_2 \leftarrow S(X^T \mathbf{u}_2, \delta(s)) / \|S(X^T \mathbf{u}_2, \delta(s))\|_2$ // soft-threshold and normalize, Algorithm 2
        $d \leftarrow \frac{(\mathbf{u}_1^T \mathbf{X} \mathbf{v}_1 + \mathbf{u}_2^T \mathbf{X} \mathbf{v}_2)}{N}$
        $score\_new \leftarrow 2d(\mathbf{u}_1^T \mathbf{X} \mathbf{v}_1 + \mathbf{u}_2^T \mathbf{X} \mathbf{v}_2) - Nd^2$
        $err \leftarrow \frac{|score\_new - score|}{|score|}$
        $rss \leftarrow \|\mathbf{X} - d(\mathbf{u}_1 \mathbf{v}_1^T + \mathbf{u}_2 \mathbf{v}_2^T)\|_F^2$
        $score \leftarrow score\_new$
**until** $interation\_count > max\_iter$ **or** $err \leq tol$
**return** $\mathbf{v}_1, \mathbf{v}_2$, $\mathbf{u}_1, \mathbf{u}_2$, $d$, $rss$

**Algorithm 2** Soft-thresholding with L1 and L2 constraints

**Input:** $\mathbf{x}$ vector
**Output:** $\mathbf{x}$ vector constrained to satisfy $l_1$ and $l_2$ constraints
**if** $\|\mathbf{x}\|_1 / \|\mathbf{x}\|_2 \leq s$ **then**
    **return** $\mathbf{x}$ // No thresholding needed
**else**
    $\tilde{\mathbf{x}} \leftarrow \mathrm{Sort}(|\mathbf{x}|, \mathrm{decreasing})$
    // Algorithm 7 from [48]
    Define $\psi(c) \leftarrow \|S(\mathbf{x}, c)\|_1 / \|S(\mathbf{x}, c)\|_2$
    Find $i$ such that $\psi(\tilde{\mathbf{x}}_i) < s < \psi(\tilde{\mathbf{x}}_{i+1})$
    Define $\delta \leftarrow \frac{\|S(x, \tilde{x}_i)\|_2}{i} \left( s\sqrt{\frac{i - \psi(\tilde{\mathbf{x}}_i)^2}{i - s^2}} - \psi(\tilde{x}_i) \right)$
    $s_{opt} \leftarrow \max(\tilde{\mathbf{x}}_i - \delta, 0)$
    **return** $S(\mathbf{x}, s_{opt})$
**end if**

#### S3 Text Performance evaluation methodology

##### A Synthetic Datasets

To assess the performance of COFE, we generated simulated datasets, each of which is an $N \times p$ matrix with a fraction $r$ of rhythmic features. The times at which the samples were collected, $t_1, \ldots, t_N$, the times COFE aims to predict, were chosen uniformly at random across one cycle of the underlying rhythm (one cycle spans $T$ units of time). To include a wide range of non-sinusoidal rhythmic patterns in the data, the feature expression at each $t_i$ was generated using periodic spline interpolation of 3 randomly chosen time-expression value pairs (see examples in S1A Fig.). Rhythmic features were standardized to have unit peak-to-trough amplitude and the arrhythmic features were assumed to be constant. All features were corrupted by additive white Gaussian noise $\mathcal{N}(0, \sigma^2)$. Thus, the SNR of the rhythmic features was $1/\sigma$. We generated synthetic datasets for combinatorial combinations of $N = 100, 200, 400$, $p = 2000$, $r = 0.1, 0.2, 0.33, 0.5$ and SNR=$0.5, 1.0, 2.0, 4.0$.

##### B Benchmark Datasets

We also analyzed the performance of COFE on time-series biological datasets (S1 Table) with the aim of comparing the predicted sample time-labels with true sample times.

##### C Metrics

We quantified two key aspects of the performance of COFE. COFE estimates the timing of the $N$ samples within one cycle. These estimates are $\hat{t}_1, \ldots, \hat{t}_N \in [0, 1)$. To determine the reordering performance, we used median absolute position error (MAPE) defined as $\min_\psi \text{median}_{i=1,\ldots,N} \{|(\hat{t}_i - t_i/T - \psi) \mod 1|\}$ [5]. MAPE always lies in [0,0.5] and error as a fraction of the rhythm period. We can only determine the relative ordering of the samples within one cycle (hence we can estimate only $t_i/T \mod 1$) and not the absolute timing $t_i$ of the samples (hence the search over different offsets $\psi$). As a rule of thumb, good performance is an MAPE less than 0.05, i.e., an error in timing reconstruction less than 5% of the rhythm period.

To quantify the ability of COFE to select exclusively rhythmic features to perform the reordering, we defined the fraction of features used for reordering that are rhythmic and the fraction of truly rhythmic biomarkers used for reordering. These are the 'precision' and 'recall' of the biomarker selection, respectively.

### S4 Text Analysis of TCGA data

#### A Data acquisition

Standardized (aligned and quantified) RNA-seq data, including from TCGA, was retrieved by means of recount3 (Bioconductor package v1.14.0)[51] annotated based on the GENCODE v29 (Ensemble 94). Unlike previous studies that attempted reordering of samples, we did not regress away any confounding variables and used raw data as is. We included only the histological types of each AC (Table S2) that included at least 250 samples. Moreover, we retained only a single tumor sample per patient per AC to train COFE (TCGA includes multiple tumor samples from some patients). We further kept protein-coding genes with an average read count of 64 reads/sample in the data. Gene expression was expressed in $\log_2$ cpm using DESeq2 (v1.40.2).

#### B COFE analysis

The primary analysis using COFE was performed using Python (S7 Table). The TCGA data was filtered to only keep genes (features) with mean $\log_2$-fold cpm of at least 7. COFE was run on all the 11 ACs individually with default parameters: 3-repeated 5-fold cross-validation with 5 restarts ($K = 5$, *repeats*=3, *restarts*=5). By default, COFE returns time-labels with respect to a normalized cycle in the interval [0, 1). We converted these time-labels to circadian time by multiplication with 24. Subsequent analyses and visualization was performed in R (S7 Table).

#### C Alignment of time-labels across ACs

The data-driven time-labels predicted by COFE are internally consistent, but the nature of the problem does not allow prediction of the absolute time-labels (the time reference is arbitrary) or the direction of time without additional information. We resolve this uncertainty by using the set of 15 expressed core clock genes to search for the consensus arrangement across all possible time references and directions of time in each AC. For this optimization, we expressed rhythmic gene expression of clock genes as phasors ($Ae^{i\phi}$) and used the complex SVD [12] to find the combination of time reference and direction for the ACs that maximized the explained variance. The arrangement that maximized this alignment was considered the consensus core clock peak time arrangement.

#### D Circadian parameters

Using the time-labels predicted by COFE ($t_n$ for each sample $n$ in the data $\mathbf{X}$), we have a complete time-series transcriptome and cancer-relevant proteome for each AC. We inferred circadian parameters from the time-series of feature $p$ by fitting a cosinor function of the form $A_p \cos(\omega(t_n - t_p))$, $\omega = 2\pi/24$ [25], where $A_p$ is the feature amplitude and $t_p$ is the peak time of expression of feature $p$.

### Supporting Information: Tables

| Tissue | Sampling | Resolution | GEO/SRA Accession | Publication |
|---|---|---|---|---|
| mouse liver | independent | 1h over 48h | SRP197108 | Pan *et al.* [52] |
| human blood monocytes | longitudinal | 3h over 42h | | Wittenbrink *et al.* [23] |
| human dermis | longitudinal | 4h over 24h | GSE205155 | del Olmo *et al.* [13] |
| human epidermis | longitudinal | 4h over 24h | GSE205155 | del Olmo *et al.* [13] |
| malaria parasite | longitudinal | 3h over 63-72h | GSE141653 | Smith *et al.* [26] |

S1 Table: Time-series biological datasets used for benchmarking COFE

| Adenocarcinoma | Histological Type |
|---|---|
| BRCA | Infiltrating Ductal Carcinoma |
| KIRC | Kidney Clear Cell Renal Carcinoma |
| PRAD | Prostate Adenocarcinoma Acinar Type |
| OV | Serous Cystadenocarcinoma |
| BLCA | Muscle invasive urothelial carcinoma (pT2 or above) |
| UCEC | Endometrioid endometrial adenocarcinoma |
| COAD | Colon Adenocarcinoma |
| THCA | Thyroid Papillary Carcinoma - Classical/usual |
| LIHC | Hepatocellular Carcinoma |
| LUAD | Lung Adenocarcinoma- Not Otherwise Specified (NOS) |
| KIRP | Kidney Papillary Renal Cell Carcinoma |

S2 Table: Histological type of each AC included in this study

S3 Table: Rhythmic genes identified by COFE, their circadian parameters and results of the enrichment analysis in each AC.

S4 Table: Rhythmic proteins and protein forms important to cancer in each AC.

S5 Table: FDA-approved, candidate and putative drugs their rhythmic targets in each AC.

| Adenocarcinoma | Best approximation error | Neglected Term | Neglected term as percentage of score |
|---|---|---|---|
| LUAD | 3874048.762 | 1262.792 | 0.033% |
| PRAD | 5371824.626 | 187.530 | 0.003% |
| LIHC | 3580022.982 | 64.330 | 0.002% |
| COAD | 4445436.789 | 1416.191 | 0.032% |
| BRCA | 9587674.848 | 15.158 | 0.000% |
| UCEC | 4899312.933 | 148.044 | 0.003% |
| OV | 4835704.272 | 68.397 | 0.001% |
| KIRP | 3407081.873 | 142.428 | 0.004% |
| BLCA | 4561640.812 | 1074.322 | 0.024% |
| THCA | 4186148.857 | -1.499 | 0.000% |
| KIRC | 6309507.411 | 1809.256 | 0.029% |

S6 Table: The neglected cross term in (2) at optimal sparsity for the different datasets.

| Language | Package name | Version |
| --- | --- | --- |
| Python | python | 3.9.15 |
| | numpy | 1.23.5 |
| | pandas | 1.5.2 |
| | scipy | 1.10.1 |
| | seaborn | 0.12.1 |
| | joblib | 1.2.0 |
| | biocthings_client | 0.2.6 |
| R | Bioconductor | 3.19 |
| | biomaRt | 2.60.1 |
| | car | 3.1-3 |
| | CircStats | 0.2-6 |
| | clusterProfiler | 4.12.6 |
| | DESeq2 | 1.44.0 |
| | dplyr | 1.1.4 |
| | effectsize | 0.8.9 |
| | ggnewscale | 0.5.0 |
| | ggpp | 0.5.8-1 |
| | ggraph | 2.2.1 |
| | ggrepel | 0.9.6 |
| | ggthemes | 5.1.0 |
| | msigdbr | 7.5.1 |
| | openxlsx | 4.2.7.1 |
| | org.Hs.eg.db | 3.19.1 |
| | patchwork | 1.3.0 |
| | PCAtools | 2.16.0 |
| | ReactomePA | 1.48.0 |
| | reactome.db | 1.88.0 |
| | recount3 | 1.14.0 |
| | reticulate | 1.39.0 |
| | SuperExactTest | 1.1.0 |
| | tidygraph | 1.3.1 |
| | tidyverse | 2.0.0 |
| | UpSetR | 1.4.0 |

S7 Table: List of software packages used in COFE and subsequent analyses

### Supporting Information: Figures

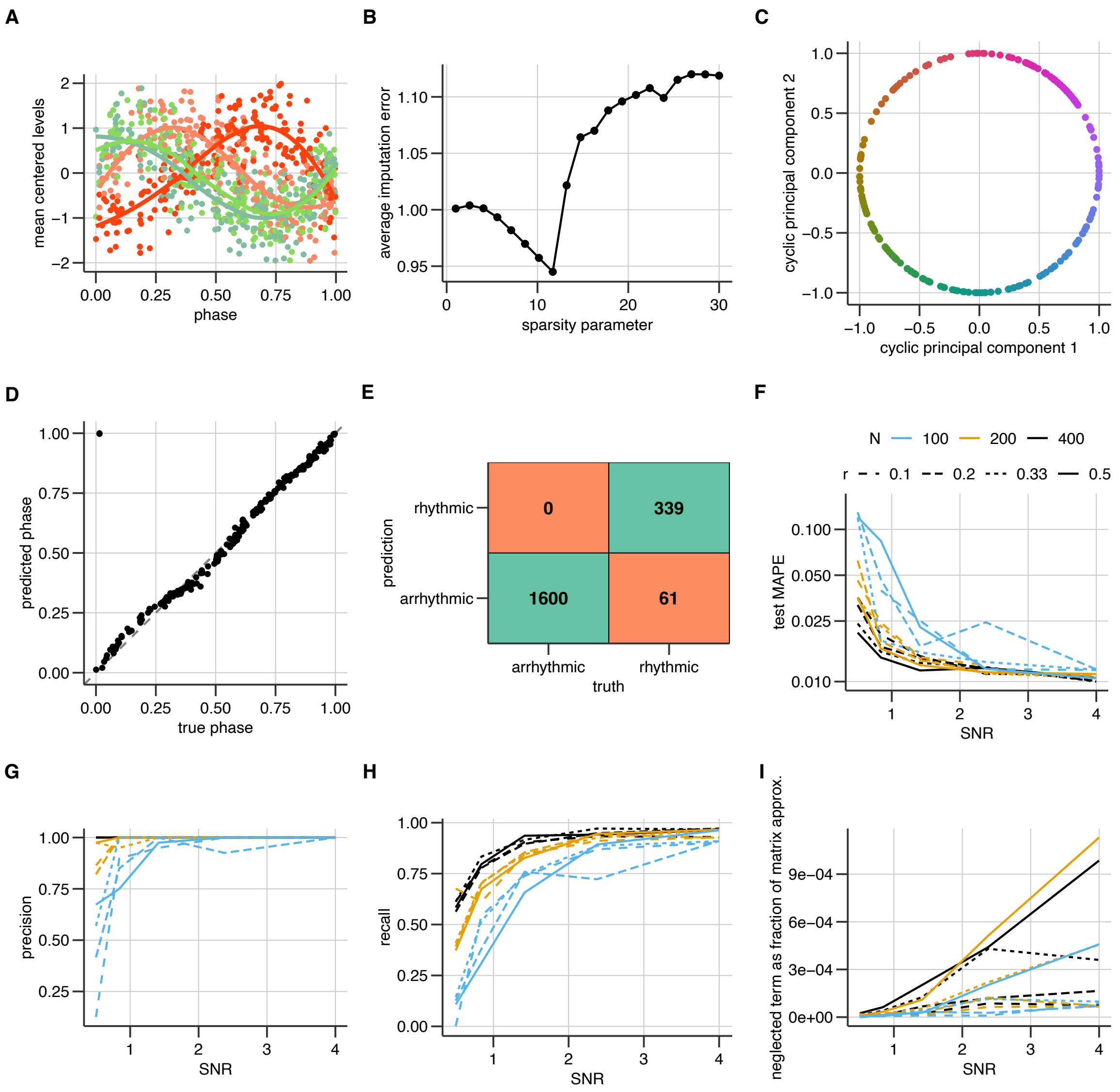


S1 Figure: **COFE benchmarking on synthetic data.** (A-E) A typical COFE run on an example data set with $N = 2000$ features of which 400 were rhythmic. (A) Raw data for a few rhythmic features (B) The output of repeated 5-fold cross-validation. (C) The underlying circular manifold reconstructed by COFE. COFE outputs the reconstructed phase within a cycle for each sample (D) and identifies rhythmic features in the data (E). (F) Temporal ordering performance of COFE measured using median absolute position error (MAPE) on data with $p = 2000$ features and different signal-to-noise ratios (SNR), number of samples ($N$) and fraction of rhythmic features ($r$). Precision (G) and recall (H) performance for rhythmic feature identification for the same synthetic data in (F). (I) The ratio of the cross term dropped from (2) as a fraction of the matrix approximation error for the synthetic data in (F).

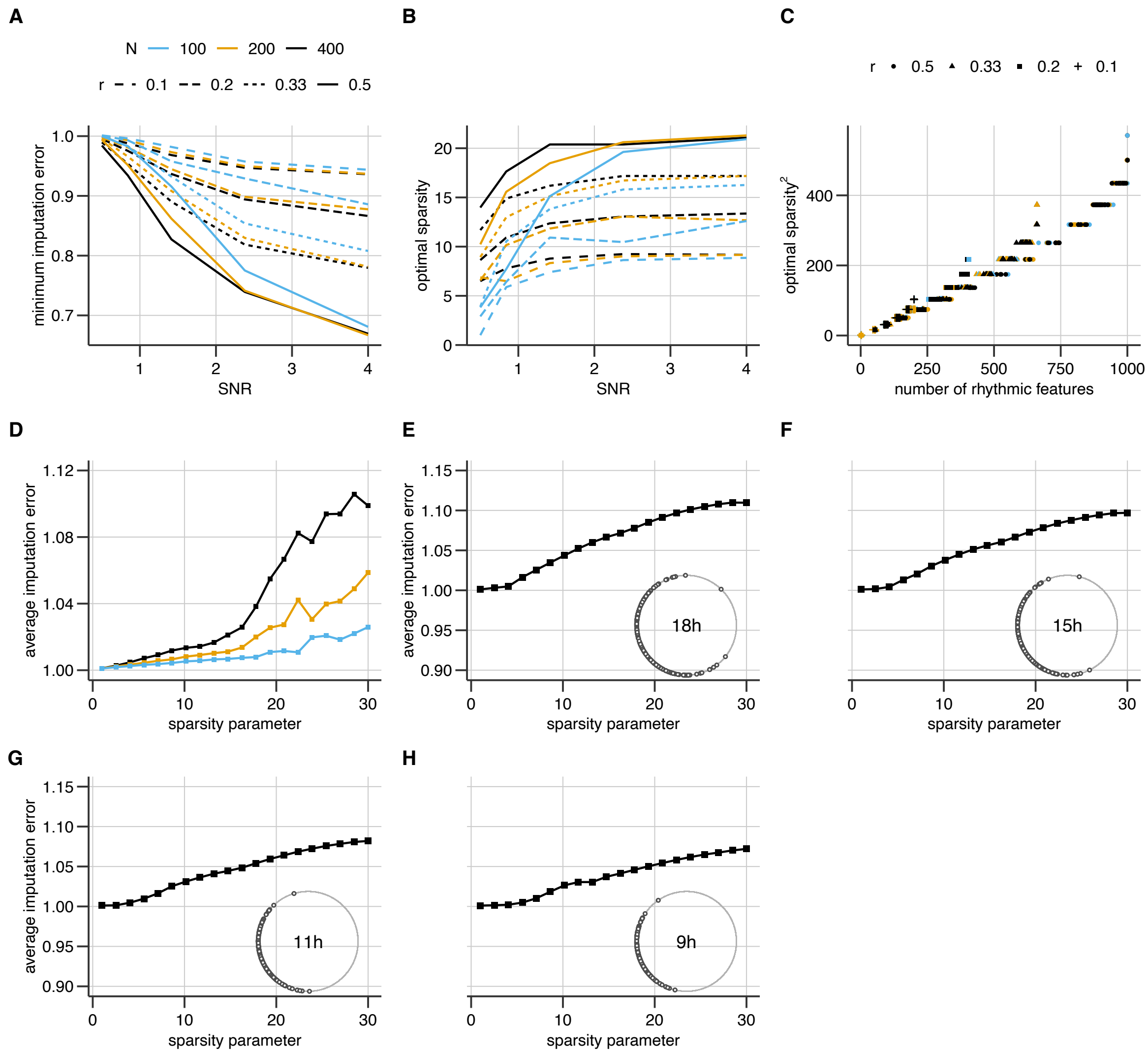


S2 Figure: **Cross-validation (CV) quantifies the quality of the ordering.** The minimum imputation error (A) and optimal sparsity parameter (B) during CV for different combinations of signal-to-noise ratios (SNR), number of samples ($N$) and fraction of rhythmic features ($r$) in (S1F Fig.). (C) The relationship between the optimal sparsity parameter and the number of rhythmic features identified. (D) Output of repeated-5-fold CV for a synthetic dataset with only arrhythmic features (2000) for different number of samples ($N$). (E-H) Output of repeated-5-fold CV on datasets with 200 samples that were restricted to different fractions of the 24h cycle (see inset), and 2000 features, of which 400 were rhythmic.

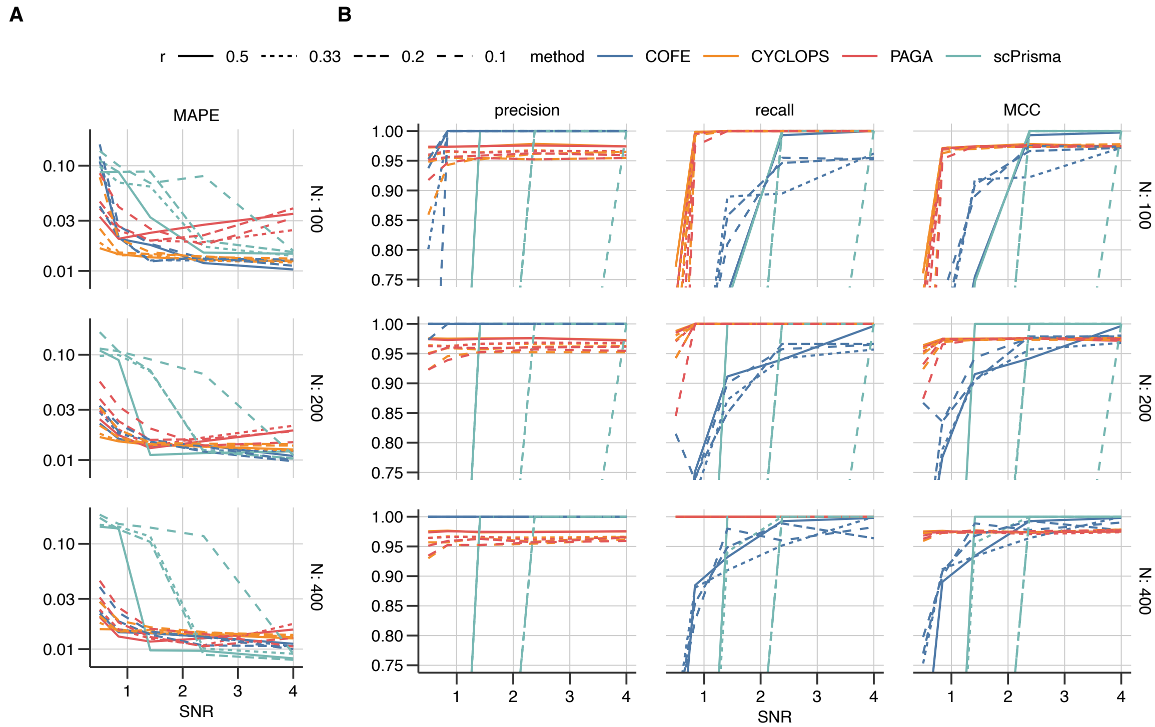


S3 Figure: **COFE benchmarking against other methods.** (A) Temporal ordering performance of COFE compared to three other methods (CYCLOPS [15], PAGA [24] and scPrisma[21]) measured using median absolute position error (MAPE) on data with $p = 2000$ features and different signal-to-noise ratios (SNR), number of samples ($N$) and fraction of rhythmic features ($r$). (B) Rhythmic feature detection performance of COFE compared to three other methods quantified by precision, recall and Matthews Correlation Coefficient (MCC), which combines precision and recall, for the same synthetic data in (A).

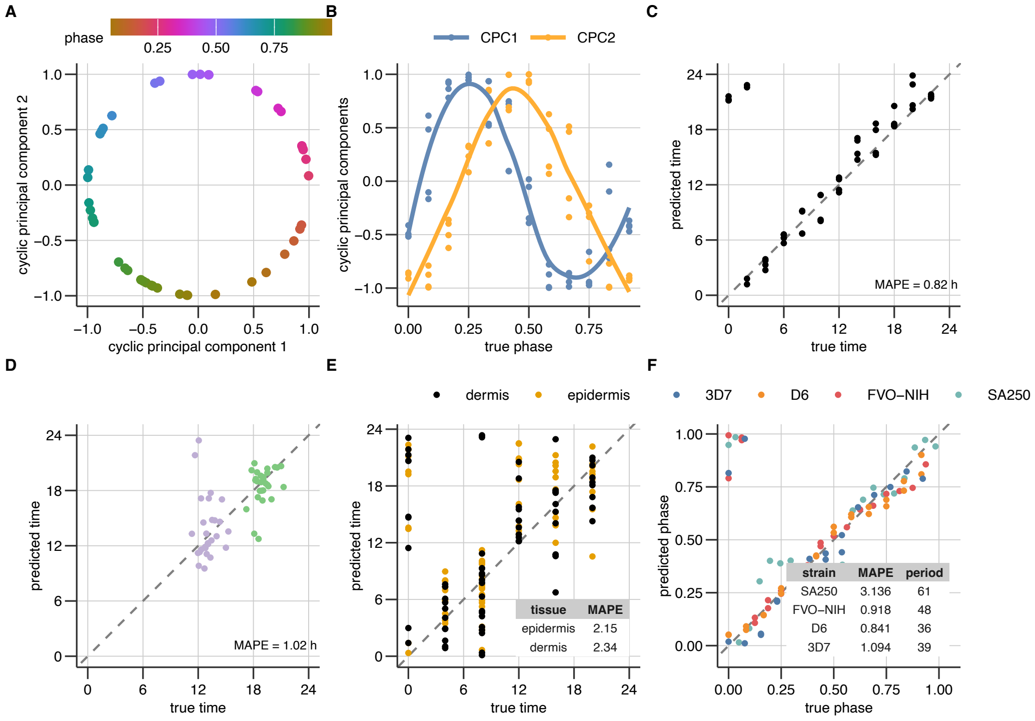


S4 Figure: **COFE benchmarking on biological data.** (A-C) COFE applied to the mouse liver RNA-seq data (SRP197108) collected hourly over 48h [52]. (A) The cyclic principal components (CPCs) extracted by COFE with the samples color-coded by phase within a cycle (normalized to 1.0). (B) The CPCs plotted against the true sample phase within a cycle (normalized to 1.0). (C) Scatter-plot of the true and predicted sample times. (D, E) Performance of COFE on human data. (D) COFE trained on Nanostring gene expression data from human blood monocytes is used to predict sample times for independent validation data [23]. (E) COFE applied to longitudinal time series human microarray gene expression data (GSE205155) from two different skin layers [13]. (F) Time label reconstruction on RNA-seq time-series of in-vitro cultures of four strains of *P. falciparum* (malaria parasite) [26].

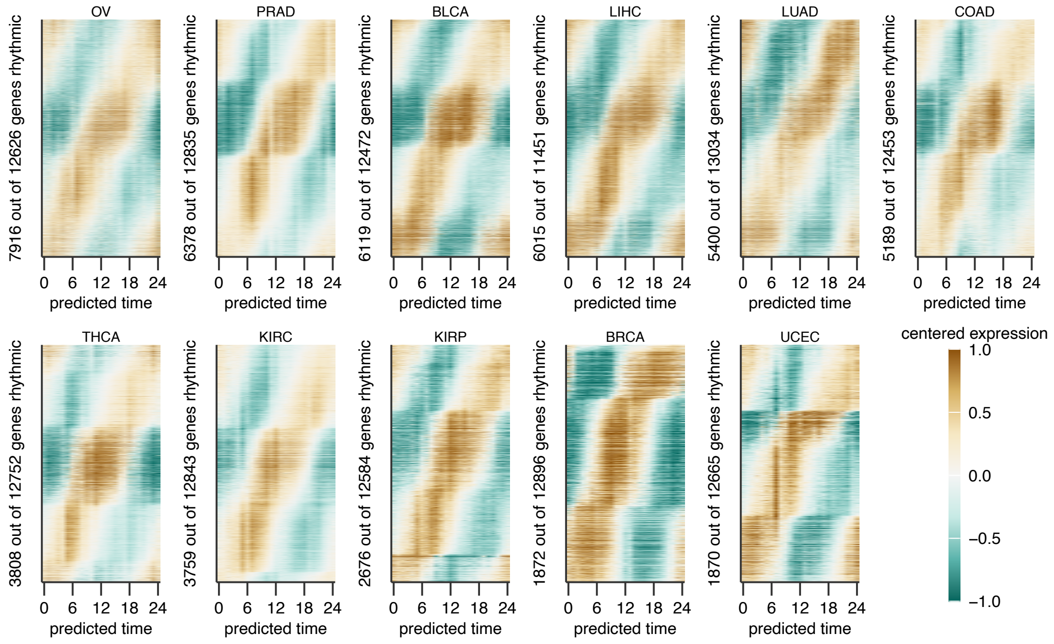


S5 Figure: **Rhythmic transcriptome in human adenocarcinomas (ACs)** Heatmaps of gene expression patterns of rhythmic genes, which are sorted by peak time of expression in each AC, for different ACs.

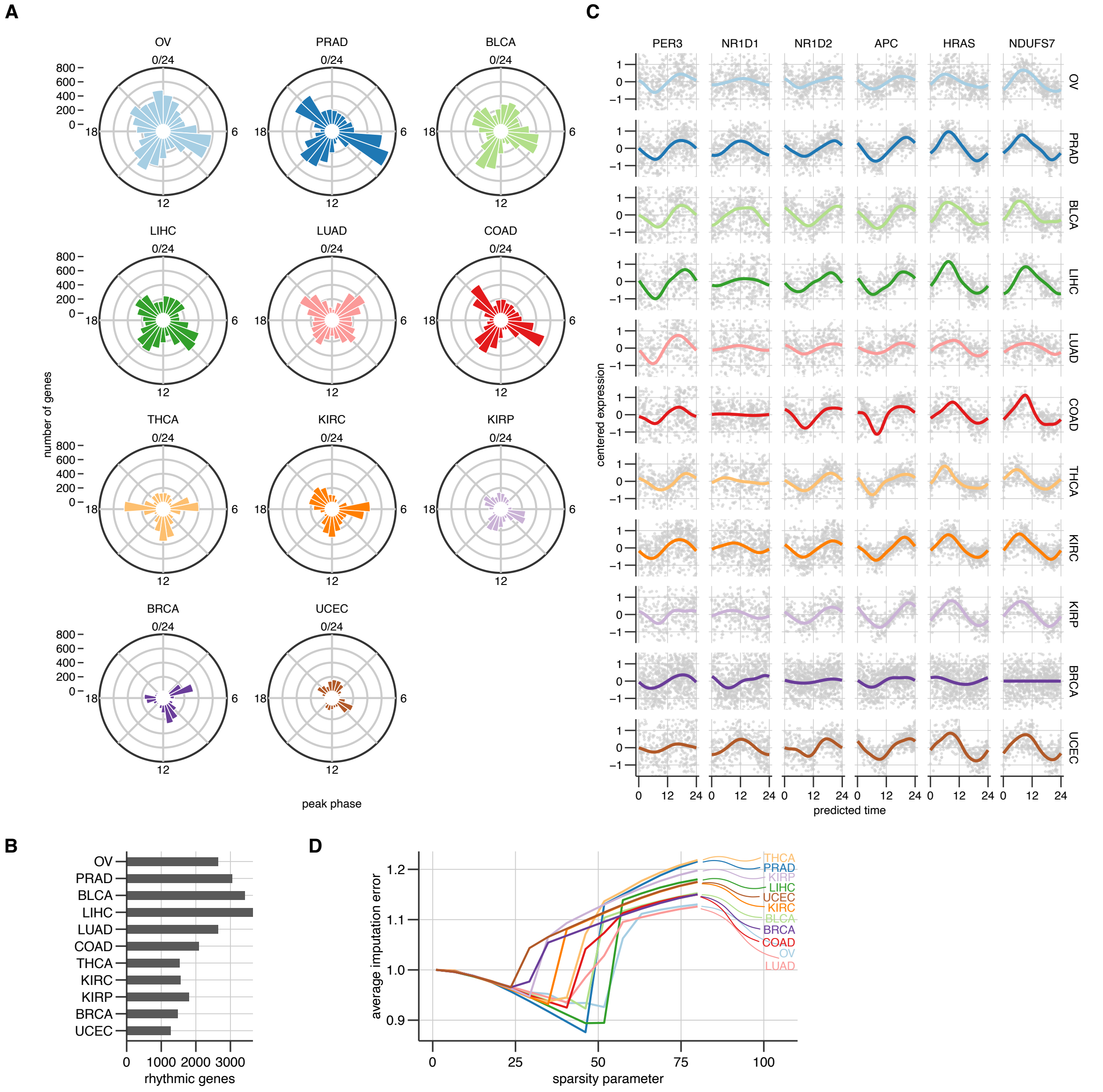


S6 Figure: **Circadian population rhythms in human adenocarcinomas (ACs).** (A) Distribution of peak phase of population rhythmic genes in different ACs. (B) The number of rhythmic genes with at least two-fold peak-to-trough amplitude in the ACs (compare with Figure 1C). (C) Raw-data ordered using predicted time-labels with the LOESS-smoothed estimates of the mean profile for selected genes. (D) Output of the repeated 5-fold cross-validation for each AC.

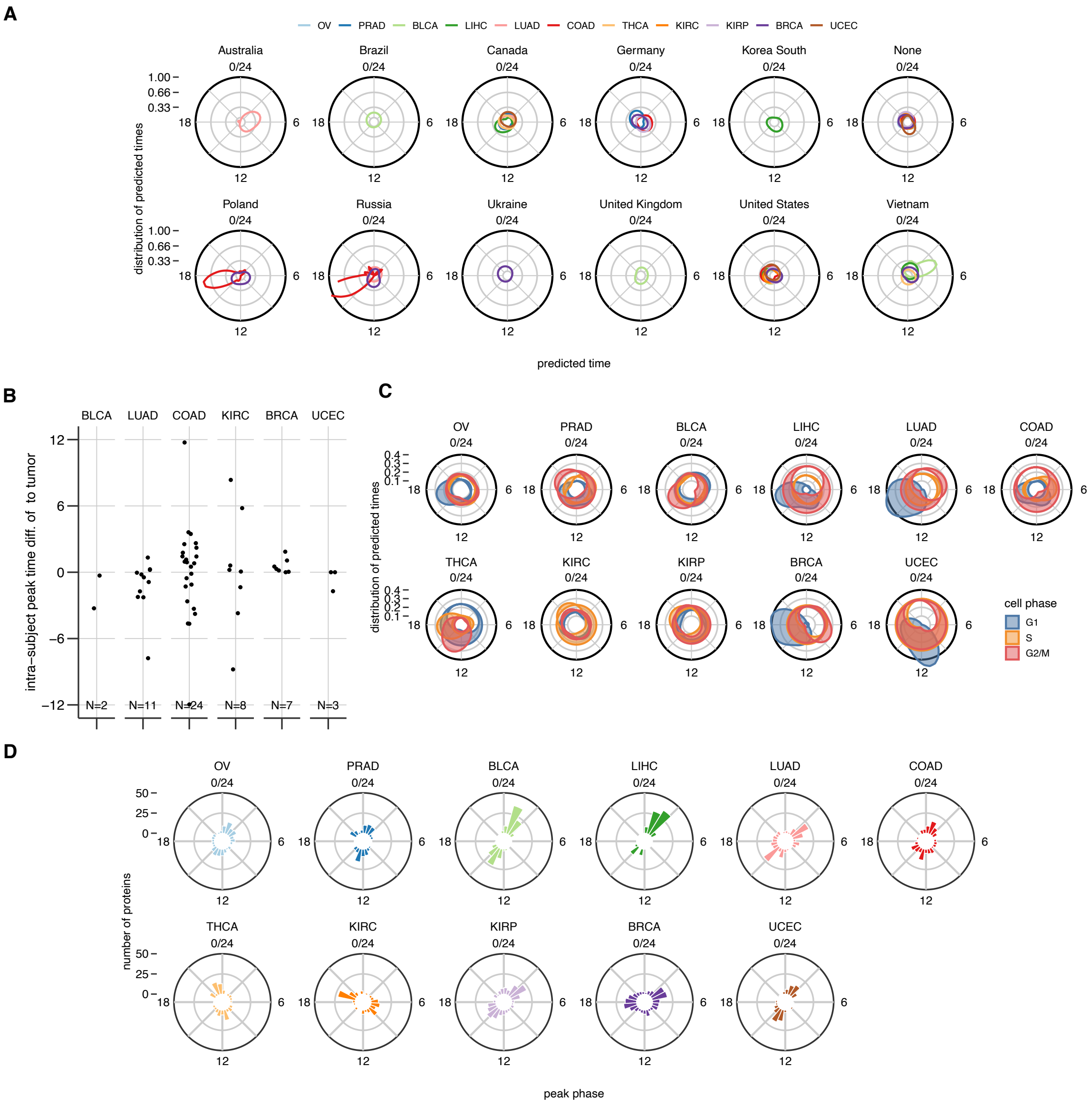


S7 Figure: **Coupling of circadian clock to the cell-cycle and the proteome.** (A) Distribution of predicted time-labels for the samples in each AC according to source country of the sample. (B) Phase difference predicted by COFE between cancer and patient-matched cancer samples not used for training COFE. (C) Predicted predominant cell-cycle phases in the different patient samples predicted by the gene expression-based cell-cycle phase scores. (D) Circular histograms of the peak time of expression of rhythmic proteins.

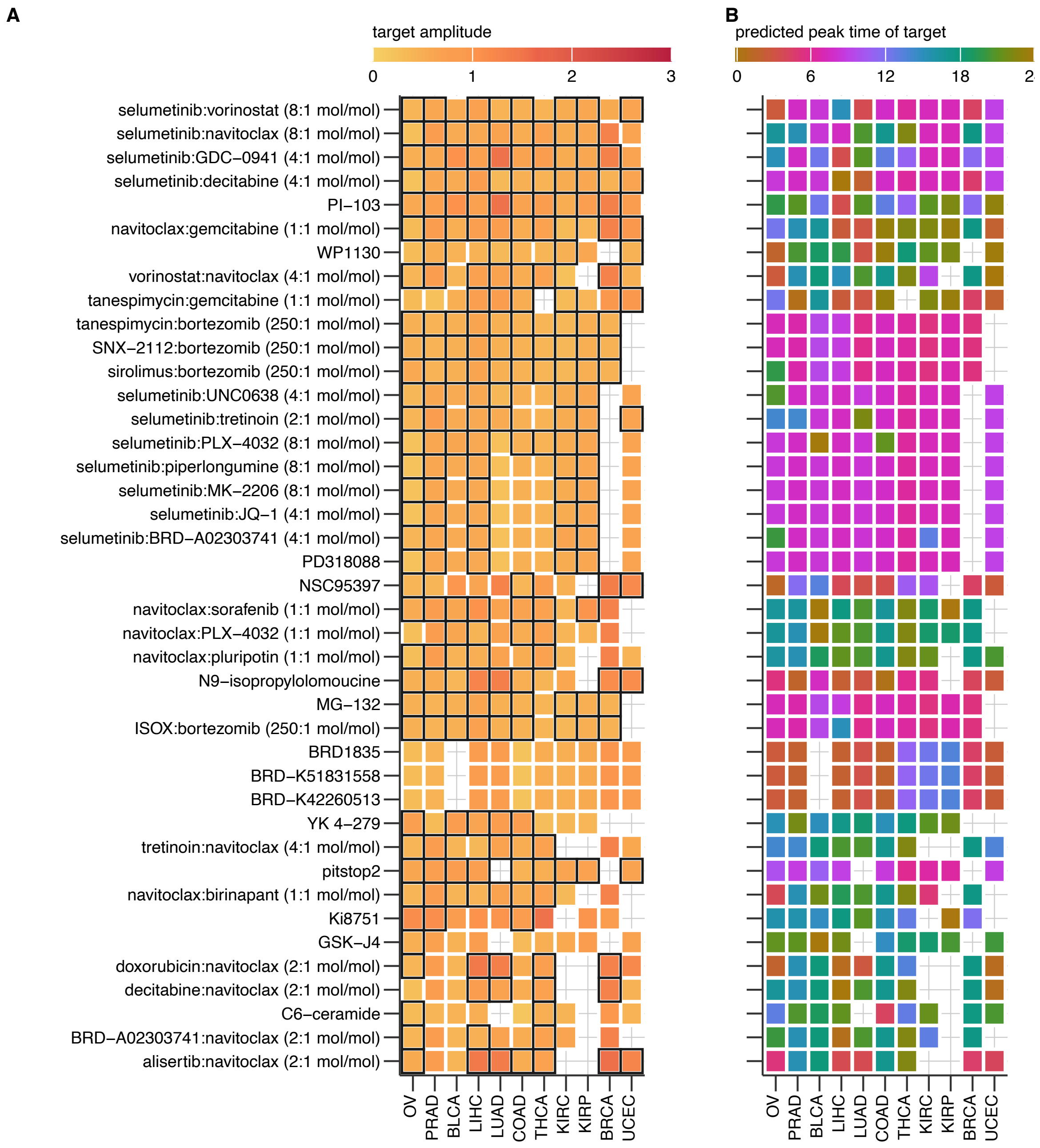


S8 Figure: **Rhythmic targets of putative cancer drugs** The $\log_2$-fold amplitude (A) and peak time relative to the core clock (B) of the largest rhythmic target gene in each AC among putative drugs that target cancer pathways or processes. All putative drugs with rhythmic targets in at least 9 ACs are included. Drugs with multiple rhythmic gene targets in an AC are boxed in black.
